# Neuronal complexity is attenuated in chronic migraine and restored by HDAC6 inhibition

**DOI:** 10.1101/2020.04.21.053272

**Authors:** Zachariah Bertels, Harinder Singh, Isaac Dripps, Kendra Siegersma, Alycia F Tipton, Wiktor Witkowski, Zoie Sheets, Pal Shah, Catherine Conway, Valentina Petukhova, Bhargava Karumudi, Pavel A. Petukhov, Serapio M. Baca, Mark M Rasenick, Amynah A Pradhan

## Abstract

Migraine is the third most prevalent disease worldwide but the mechanisms that underlie migraine chronicity are poorly understood. Cytoskeletal flexibility is fundamental to neuronal-plasticity and is dependent on dynamic microtubules. Histone-deacetylase-6 (HDAC6) decreases microtubule dynamics by deacetylating its primary substrate, α-tubulin. We use validated models of migraine to show that HDAC6-inhibition is a promising migraine treatment and reveal an undiscovered cytoarchitectural basis for migraine chronicity. The human migraine trigger, nitroglycerin, produced chronic migraine-associated pain and decreased neurite growth in headache-processing regions, which were reversed by HDAC6 inhibition. Cortical spreading depression (CSD), a physiological correlate of migraine aura, also decreased cortical neurite growth, while HDAC6-inhibitor restored neuronal complexity and decreased CSD. Importantly, a calcitonin gene-related peptide receptor antagonist also restored blunted neuronal complexity induced by nitroglycerin. Our results demonstrate that disruptions in neuronal cytoarchitecture are a feature of chronic migraine, and effective migraine therapies might include agents that restore microtubule/neuronal plasticity.

## Introduction

Migraine is an extremely common neurological disorder that is estimated to affect 14% of the world population, making it the third most prevalent disease worldwide [1, 2]. One particularly debilitating subset of migraine patients are those with chronic migraine, which is defined as having more than 15 headache days a month [3]. Despite its high prevalence, migraine therapies are often only partially effective or are poorly tolerated, creating a need for better pharmacotherapies [4]. Recent clinical success of antibodies against calcitonin gene related peptide (CGRP) and CGRP receptor demonstrate the effectiveness of targeted migraine therapeutics. While there has been more research into understanding the molecular mechanisms of migraine, there remains much to be discovered.

Neuroplastic changes play an important role in a variety of chronic neuropsychiatric conditions [5], and epigenetic alterations through histone deacetylases (HDACs) are frequently investigated. HDACs are best characterized for their ability to deacetylate histones, promoting chromatin condensation and altered gene expression [6]. Intriguingly, some HDACs can also deacetylate non-histone targets, including proteins involved in the regulation of cytoarchitecture. Due to its cytoplasmic retention signal, HDAC6 is primarily expressed in the cytosol [7], and one of its primary targets for deacetylation is α-tubulin [7, 8]. α- and β-tubulin form heterodimers that make up microtubules, which are a major component of the cytoskeleton, and regulate intracellular transport, cell morphology, motility, and organelle distribution [9, 10]. Microtubules undergo multiple cycles of polymerization and depolymerization, thus rendering them in a constant state of dynamic instability [9]. Tubulin displays a variety of post-translational modifications, including α-tubulin acetylation, which occurs endogenously through α-tubulin N-acetyltransferase I (αTAT1) and it is correspondingly deacetylated by HDAC6 [10]. Tubulin acetylation is associated with increased flexibility and stability of microtubules [11]. In contrast, deacetylated microtubules are more fragile and prone to breakage [11]. Microtubules are important for cellular response to injury and play a role in neurite branching [12]; and microtubule dynamics influence neuronal signaling and mediate axonal transport [13-17]. Importantly, changes in cellular structure such as alterations in dendritic spine density, have been implicated in disease chronicity [18-21].

The aim of this study was to determine if altered neuronal cytoarchitecture facilitates the chronic migraine state. We observed decreased neuronal complexity in headache-processing brain regions in the nitroglycerin (NTG) model of chronic migraine-associated pain. We further demonstrated that treatment with HDAC6 inhibitor reversed these cytoarchitectural changes and correspondingly decreased cephalic allodynia. These studies were extended to a mechanistically distinct model of migraine, cortical spreading depression (CSD), which is thought to be the electrophysiological correlate of migraine aura. Again, we observed decreased neuronal complexity in migraine related sites, which was reversed by HDAC6 inhibitor. To investigate the translational implication, we also tested the effect of olcegepant, a CGRP receptor inhibitor, and found that it alleviated chronic allodynia induced by NTG, and restored cytoarchitectural changes associated with chronic migraine-associated pain. These results suggest a novel mechanism for migraine pathophysiology and establish HDAC6 as a novel therapeutic target for this disorder.

## Results

### Exposure to chronic NTG induces cytoarchitectural changes in key pain processing regions

Changes in the structural plasticity of neurons have been observed in a number of neuropsychiatric disorders and can serve as a marker of disease chronicity [20, 21]. To investigate if this was also the case in migraine, we treated male and female C57BL6J mice every other day for 9 days with NTG or vehicle and tested for mechanical periorbital responses on days 1, 5 and 9 (Figure 1A). NTG induced severe and sustained cephalic allodynia as measured by von Frey hair stimulation of the periorbital region as compared to vehicle animals on the same day (Figure 1B). Mice were sacrificed on day 10, 24 h after the final NTG/VEH treatment, and neuronal size and arborization were examined through a Golgi staining procedure in a key cephalic pain processing region, the trigeminal nucleus caudalis (TNC) (Figure 1C) [22]. We observed a dramatic decrease in neuronal complexity after NTG treatment (Figure 1D). Neurons of chronic NTG-treated mice had significantly fewer branch points (Figure 1E), and shorter neurites resulting in decreased overall length of the neurons (Figure 1F). Further examination of the complexity of the neurons using Scholl Analysis, showed a significant decrease in the number of intersections following NTG treatment (Figure 1 G-I). In addition to the TNC we also determined if other brain regions related to central pain processing were affected by chronic NTG treatment. We examined the somatosensory cortex (SCx) and periaqueductal grey (PAG) of these mice and found similar results, where neurons from NTG-treated mice had fewer branch points, were shorter in length, and had fewer intersections. To ensure that this effect was associated with migraine-pain processing and not a non-specific effect of NTG we also analyzed neuronal complexity in the nucleus accumbens shell (NAc), a region more commonly associated with reward, and found no alteration in number of branches, total neuron length, or Sholl analysis for cells in this region (Supplementary Figure 1). Furthermore, we also examined the dorsal horn of the lumbar spinal cord, an important site for peripheral but not head-pain processing; and found no differences in NTG versus vehicle controls in number of branches, total neuron length, or Sholl analysis (Supplementary Figure 1). These results suggest that decreased neuronal complexity may be a feature that maintains the chronic migraine state; a previously undiscovered phenomenon.

**Figure 1.**
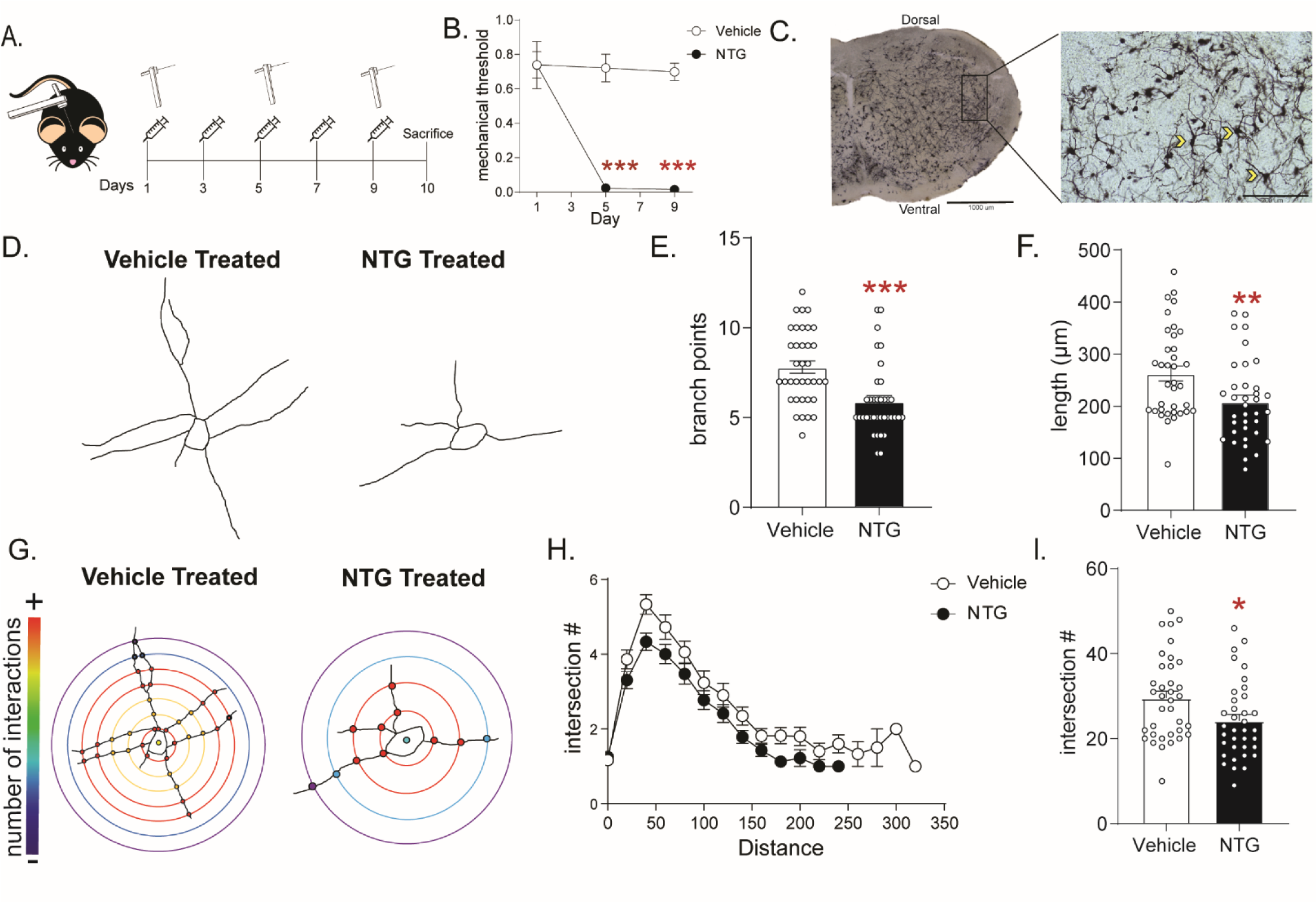
The NTG model of chronic migraine produces cytoarchitectural changes in a cephalic pain processing region. (A) Schematic of testing schedule, M&F C57Bl6/J mice were treated with vehicle or nitroglycerin (10 mg/kg, IP; NTG) every other day for 9 days. (B) Periorbital mechanical thresholds were accessed prior to Vehicle/NTG administration on days 1, 5 and 9. NTG produced severe cephalic allodynia p<0.001 effect of drug, time, and interaction, two-way RM ANOVA and Holm-Sidak post hoc analysis. ***p<0.001 relative to vehicle on day 1 n=8/group. (C) Representative image taken of Golgi stained TNC at 4x (left) and 20x (right). Chevrons indicate neurons traced from the selected image and demonstrate type of neurons selected from this region. (D) Representative tracing of neurons from mice treated with chronic vehicle (left) or NTG (right) demonstrating reduced neural processes after chronic NTG treatment. (E) The number of branch points/neuron was decreased following chronic NTG. Unpaired t-test. ***p<0.001 (F) Total neuron length was also compromised after NTG treatment. Unpaired t-test. **p<0.01. (G) Representative Sholl plots of vehicle (left) and NTG (right) treated mice. (H) Sholl analysis broken up by 20 voxel distances from the center of the cell showing differences between groups. (I) Sholl analysis revealed a significant decrease in total intersections after chronic NTG treatment. Unpaired t-test. *p<.05 n=6 mice/group, 6 neurons per mouse.

### HDAC6 inhibition increases acetylated α-tubulin and neuronal cytoarchitectural complexity

We hypothesized that if we could restore migraine-compromised cytoarchitectural complexity, that this might also relieve cephalic pain. Recent studies indicate that increased tubulin acetylation facilitates microtubule flexibility and prevents breakage [10, 23, 24]. Thus, we hypothesized that inhibiting HDAC6 to promote microtubule stability may restore the neuronal complexity observed following chronic NTG. We tested the selective HDAC6 inhibitor, ACY-738, in the chronic NTG model. Mice were treated chronically with NTG or VEH for 9 days. On day 10, mice were injected with ACY-738, after 4h, tissue was processed and analyzed by western blot. ACY-738 treatment resulted in a significant increase in the ratio of acetylated α-tubulin to total tubulin in three key migraine processing regions; the trigeminal ganglia (TG), TNC, and SCx (Supplementary Figure 2). A separate group of mice were analyzed by neuronal tracing in the TNC, 4h post-ACY-738 (Figure 2A). Again, chronic NTG treatment caused a decrease in branch points (Figure 2B), combined neurite length (Figure 2C), and number of intersections (Figure 2D-F). In contrast, treatment with ACY-738 led to a significant increase in these measures in both chronic vehicle and NTG groups (Figure 2A-F).

**Figure 2.**
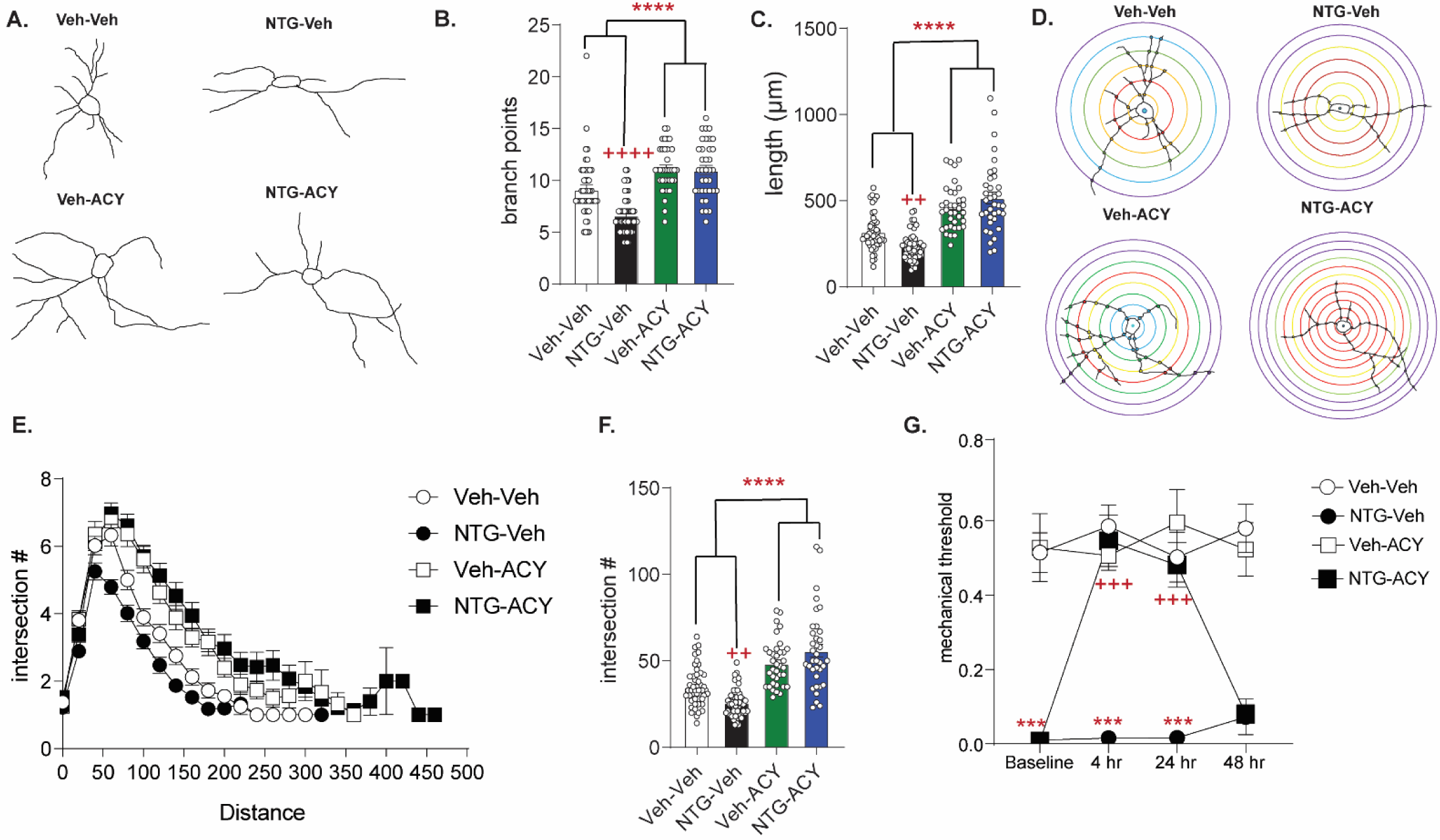
Treatment with HDAC6 inhibitor restores blunted neuronal complexity and inhibits migraine-associated pain. (A) Representative neuron tracing of mice that were chronically treated with vehicle or NTG (10 mg/kg, IP) every other day for 9 days and on day 10 were injected with ACY-738 (50 mg/kg, IP) or vehicle (5 % DMSO in 0.9% NaCl, IP) and sacrificed 4h later for Golgi staining. (B) The number of branch points/neuron were significantly decreased following NTG treatment (NTG-Vehicle); while ACY-738 treated mice showed increased branching irrespective of pretreatment. p<0.0001 effect of chronic treatment, drug treatment, and interaction, two-way ANOVA and Holm-Sidak post hoc analysis. ****p<0.001 effect of ACY-738; ++++p<.001 effect of NTG. (C) NTG also decreased total pixel length/neuron, while ACY-738 treatment increased it. p<0.0001 effect of NTG treatment, drug treatment, and interaction, two-way ANOVA and Holm-Sidak post hoc analysis. ****p<0.0001 effect of ACY-738; ++p<0.01 effect of NTG. (D) Representative Sholl analysis image of neuronal complexity in the four groups. (E) Sholl analysis broken up by 20 voxel distances reveal differences between NTG and ACY-738 treatment. (F) Chronic NTG results in significantly fewer total interactions relative to vehicle treatment. ACY-738 increases total interactions in both vehicle and NTG groups. p<0.0001 effect chronic treatment, drug treatment, and interaction, two-way ANOVA and Holm-Sidak post hoc analysis. ****p<0.001 effect of ACY-738; ++p<0.01 effect of NTG. For all analysis n=6 mice/group, 6-8 neurons per mouse. (G) C57Bl6/J mice underwent chronic intermittent NTG/Veh treatment for 9 days, on day 10 basal mechanical thresholds were assessed, and mice were subsequently injected with ACY-738 (50 mg/kg IP) or Vehicle and tested 4, 24, or 48 h later. Chronic NTG treatment caused severe cephalic allodynia (Baselines); which was significantly inhibited by ACY-738 at 4h and 24h post-injection. p<.001 drug, time, and interaction, two-way RM ANOVA, Holm-Sidak post hoc analysis, ***p<.001 as compared to Veh-Veh treated mice; +++p<.001 as compared to the NTG-Veh treated mice; n=12 mice/group.

### HDAC6 inhibition reverses NTG-induced allodynia

Next, we determined if this restored neuronal complexity would affect behavioral outcomes. Mice were treated with chronic intermittent NTG or vehicle for 9 days. On day 10, baseline cephalic allodynia was observed in mice treated chronically with NTG but not vehicle (Figure 2G, baselines). Mice were then injected with ACY-738 or vehicle and tested 4, 24, or 48h later. ACY-738 significantly reversed cephalic allodynia in NTG treated mice for up to 24h post-injection. Mechanical responses in vehicle-ACY-738 treated animals were unaffected (Figure 2G), which suggests that HDAC6-augmented neuronal complexity in a pain-free animal does not alter endogenous pain processing. Interestingly, the half-life of ACY-738 is only 12 minutes [25], thus short-term inhibition of HDAC6 still produced long-lasting behavioral and cytoarchitectural changes.

We confirmed that this behavioral effect was due specifically to changes in HDAC6 inhibition. We first tested two pan-HDAC inhibitors: the well characterized inhibitor trichostatin A (TSA, Figure 3A); and a novel brain-penetrant pan-HDAC inhibitor, RN-73 [26] (Figure 3B). Both significantly reversed chronic NTG-induced allodynia, albeit for a much shorter duration than ACY-738. In contrast, when we tested the Class I, HDAC1 and 2 selective inhibitor, ASV-85 (Extended Table 3.1), we did not observe any change in NTG-induced chronic allodynia relative to vehicle controls (Figure 3C). These data further support our finding that chronic migraine-associated pain can be blocked, specifically, by HDCA6 inhibition, and that this effect is due to increased acetylation of cytosolic HDAC6 substrates, rather than histone acetylation in the cell nucleus.

**Figure 3.**
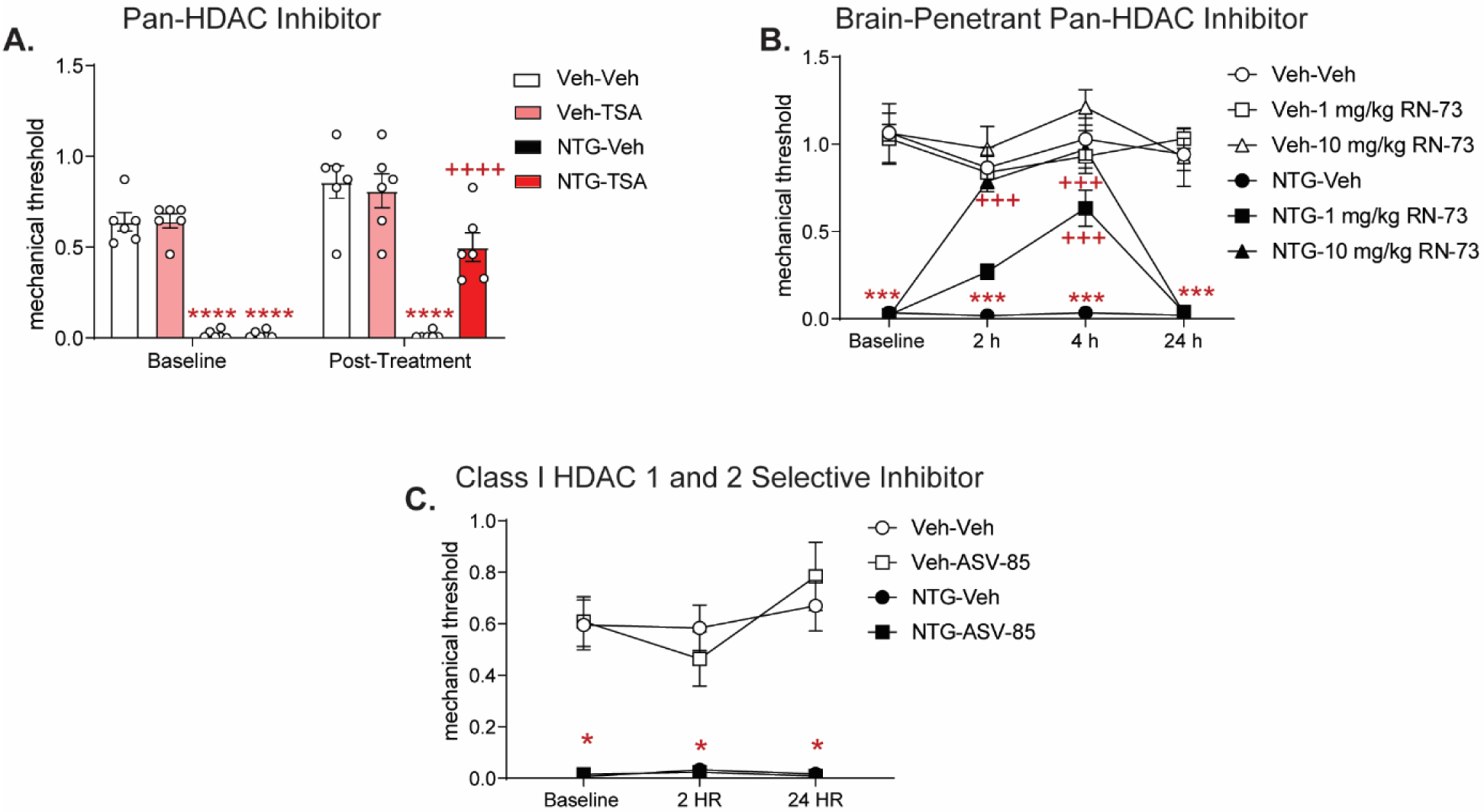
Pan-HDAC inhibitors, but not Class I-selective HDAC inhibitor block chronic migraine-associated pain. Male and female C57BL6J mice were treated with chronic intermittent NTG (10 mg/kg IP) or Vehicle for 9 days. On day 10 mice were subsequently tested for baseline responses (Baseline) and then injected with various HDAC inhibitors. Baselines were always lower for NTG-treated mice demonstrating chronic cephalic allodynia. Separate groups of mice were tested for each drug. (A) Mice were treated with the pan-HDAC6 inhibitor Trichostatin A (TSA, 2 mg/kg IP) or Vehicle (20% DMSO in 0.01 M PBS IP) and subsequently tested 2 h later. TSA significantly inhibited chronic cephalic allodynia. p<0.05 effect of treatment, drug, time, and interaction, Three-way ANOVA and Holm-Sidak post hoc analysis. ****p<0.0001 relative to Vehicle-Vehicle, ++++p<0.0001 relative to NTG-Vehicle at same time point n=6-10/group (B) Mice were treated with the novel brain-penetrant pan-HDAC inhibitor, RN-73 at 1 or 10 mg/kg (IP) or vehicle (10% DMSO, 10%Tween-80, and 0.9% NaCl). Mice were subsequently tested in the hind-paw region 2, 4, and 24 h post treatment. RN-73 at the 10 mg/kg dose had a significant effect at the 2 and 4 h time point, and the 1 mg/kg dose had a significant effect only at the 4 h time point compared to the NTG-Veh group. Neither dose of RN-73 produced any effect 24 h after treatment compared to NTG-Veh. Three-way RM ANOVA and Holm-Sidak post hoc analysis, p<0.05 effect of treatment, drug, time and interaction, ****p<0.001 relative to Vehicle-Vehicle at matching time point, +++p<.001 NTG-RN-73 relative to NTG-Vehicle at same time point, n=8/group. (C) Following chronic NTG/VEH treatment, mice were treated with the Class I specific HDAC inhibitor, ASV-85 (1 mg/kg IP) or Vehicle (6.25% DMSO, 5.625% Tween-80, and 0.9% NaCl, IP) and had subsequent mechanical thresholds taken at 2 and 24 h post-treatment. ASV-85 failed to inhibit NTG-induced pain. NTG-ASV-85 and NTG-Veh treated mice both were significantly different than the Vehicle control groups at both 2 and 24 h time points, Three-way ANOVA and Turkey post hoc analysis p<.05 effect of treatment, *p<.05 NTG treated mice compared to Vehicle treated mice at same time point.

### HDAC6 mRNA and protein is found ubiquitously in key migraine processing regions

HDAC6 expression is enriched in certain brain regions, such as the dorsal raphe [27] and, to the best of our knowledge, HDAC6 expression in head pain processing regions is not well characterized. In situ hybridization using RNAScope and immunohistochemical analysis (Supplementary Figure 3A,B) revealed abundant expression of HDAC6 transcripts in TG, TNC, and SCx. Gene expression analysis revealed that, of these regions, chronic NTG treatment increased HDAC6 expression in the TG (Supplementary Figure 3C), which are the first order cells regulating cephalic pain processing. Thus, HDAC6 is expressed and regulated dynamically in regions that are critical for migraine-associated pain processing.

### HDAC6 inhibitor results in reduced CSD events

CSD is an electrophysiological property thought to underlie migraine aura. It is mechanistically and etiologically distinct from the NTG model of migraine pain, and reduction of CSD events is a feature of many migraine preventives [28]. Thus, we examined whether CSD propagation was also affected by HDAC6 inhibition. Briefly, the skull was thinned in an anesthetized animal to reveal the dural vasculature and cortex underneath (Figure 4A). Two burr holes were made, and the more rostral was used to continuously drip KCl onto the dura to induce CSD, while local field potentials (LFPs) were recorded from the caudal burr hole. The somatosensory/barrel cortex was targeted, as it is more sensitive to CSD induction [29]. Throughout the 1h recording, CSDs were identified by visual shifts in light and sharp decreases in the LFP (Figure 4B-C). Pretreatment with ACY-738 resulted in significantly fewer CSD events relative to vehicle controls (Figure 4D), indicating that HDAC6 inhibition also effectively blocks this separate migraine mechanism.

**Figure 4.**
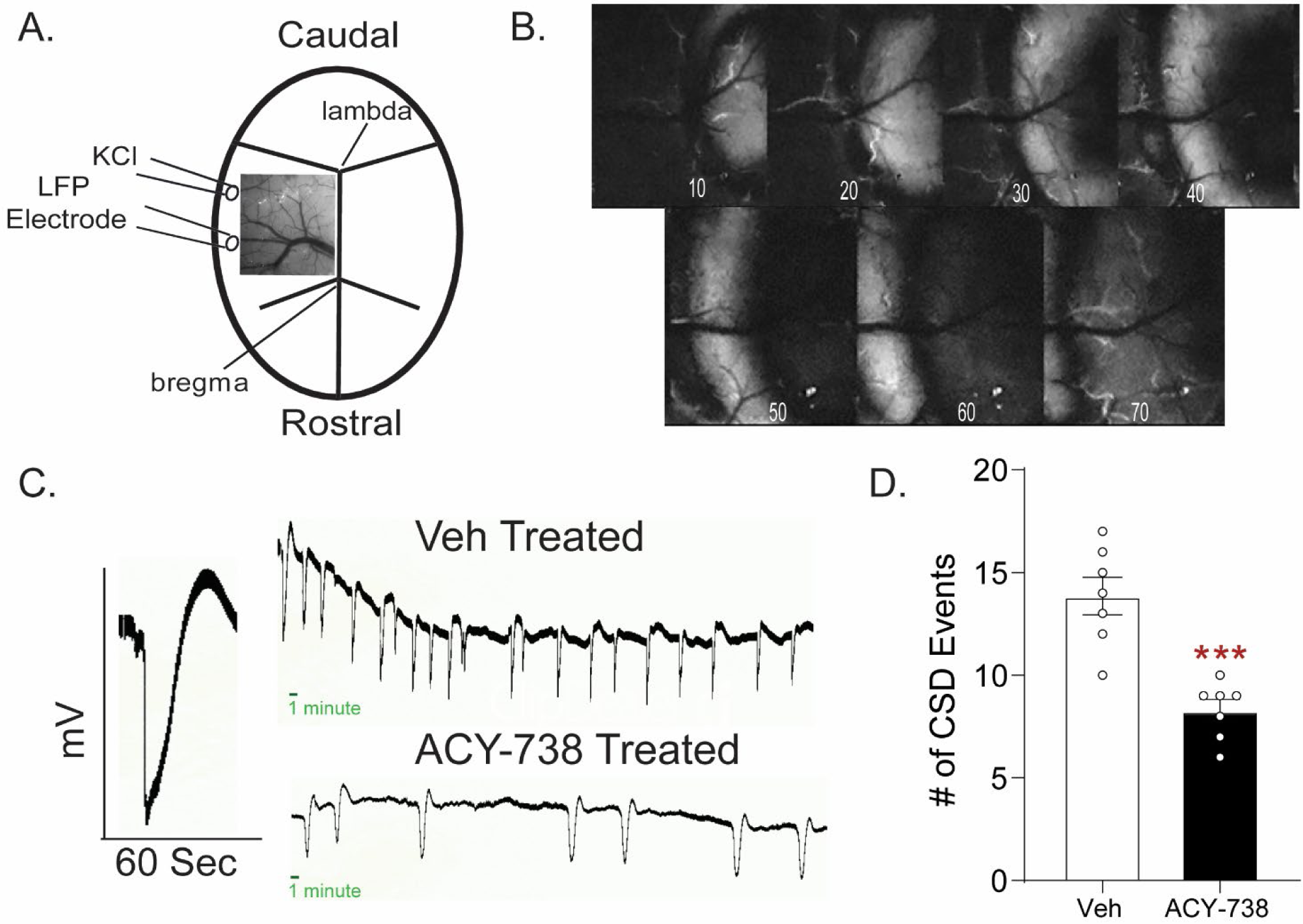
ACY-738 reduces cortical spreading depression events. (A) Schematic of the thinned skull preparation used to visualize CSD and placement of KCl infusion and LFP recording. (B) Image sequence shows the wave of change in reflectance associated with a CSD event. (C) Representative tracing of a single CSD event of voltage change versus time. Representative line tracing of CSDs in a Vehicle (Top) vs. ACY-738 (Bottom) treated mouse over a 1h period. (D) Animals pretreated with ACY-738 (50 mg/kg IP) 4h before CSD recordings began showed a significant reduction in the average number of CSD events recorded over an hour. Unpaired t-test. ***p<0.001 n=7/group.

### CSD results in decreased neuronal complexity in the somatosensory cortex that is restored by HDAC6 inhibition

We next examined the neuronal complexity of pyramidal neurons within the somatosensory cortex following CSD induction. Sham mice that underwent anesthesia and surgery, but did not receive KCl, were used as controls. Mice were pretreated with ACY-738 or vehicle, underwent CSD or sham procedure, and were immediately sacrificed for Golgi staining of the SCx (Figure 5 A-B). In the somatosensory cortex, CSD evoked a significant decrease in branch points (Figure 5C) and total length of neurons (Figure 5D). In addition, CSD also resulted in a significant reduction in the number of branches in neurons of the TNC (Supplemental Figure 4), a region that is known to be activated following CSD events [30]. In contrast, ACY-738 increased neuronal complexity in the cortex in both sham and CSD groups. Sholl analysis demonstrated a dramatic decrease in neuronal complexity after CSD, while ACY-738 treatment had the opposite effect (Figure 5 E-G). These results demonstrate that decreased neuronal complexity is also observed in a second, mechanistically distinct model of migraine, and that HDAC6 inhibition can prevent these changes in neuronal cytoarchitecture and decrease CSD events.

**Figure 5.**
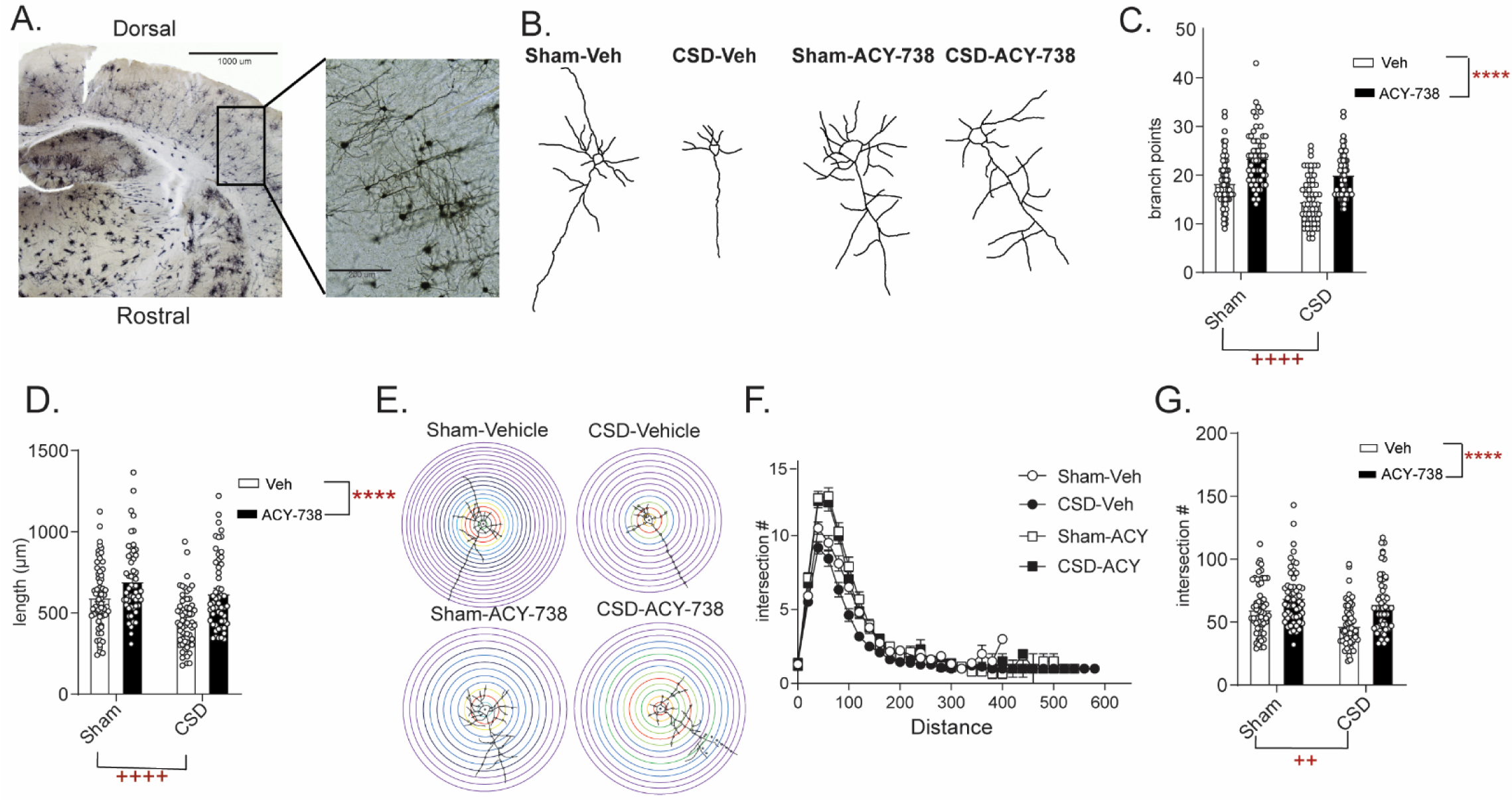
CSD induces decreased neuronal complexity that is prevented by treatment with ACY-738. (A) Representative image of Golgi stained sensory/barrel cortex at 4x (left) and 20x (right). (B) Representative neuronal tracing for mice that underwent pretreatment with Vehicle or ACY-738 and underwent Sham or CSD procedures. (C) Analysis of number of branch points/neuron reveal a significant effect of CSD and of ACY-738. Two-way ANOVA with Holm-Sidak post hoc analysis. ++++p<0.0001 effect of CSD; ****p<0.0001 effect of ACY-738. (D) Neurons were further analyzed for pixel length per neuron and CSD significantly decreased overall length, while ACY-738 significantly increased length. Two-way ANOVA and Holm-Sidak post hoc analysis. ++++ p<0.0001 effect of CSD, **** p<0.0001 effect of ACY-738 (E) Representative Sholl analysis plot of a neuron demonstrating CSD reduces and ACY-738 increases neuronal complexity. (F) Sholl analysis broken up by 20 voxel distances showing differences between groups. (G) Sholl analysis revealed a significant decrease in total intersections after CSD compared to Sham mice; and pretreatment with ACY-738 increased total intersections compared to vehicle treated groups. Two-way ANOVA and Holm-Sidak post hoc analysis ++p<0.01 effect of CSD, **** p<0.0001 effect of ACY-738. n=6 mice/group, 9 neurons/mouse

### CGRP receptor blockade reverses NTG-induced chronic allodynia and cytoarchitectural alterations

We next sought to determine if migraine-selective therapies could influence neuronal cytoarchitecture; and we tested the small molecule CGRP receptor antagonist, olcegepant, in the chronic NTG model [31]. Mice developed a sustained allodynia to repeated NTG treatment (Figure 6A). On day 10, 24 hours after the final NTG injection, baseline mechanical responses were assessed and mice were treated with olcegepant or vehicle. Olcegepant significantly inhibited NTG-induced cephalic allodynia (Figure 6B), similar to previously published reports [32]. Subsequent golgi analysis of TNC revealed cytoarchitectural alterations in this cohort of animals (Figure 6C). As was observed previously, chronic NTG treatment decreased the number of branch points (Figure 6D), combined neurite length (Figure 6E), and number of intersections using Sholl analysis (Figure 6F-H). Interestingly, olcegepant treatment restored neuronal complexity induced by chronic NTG, but had no effect in chronic vehicle treated mice (Figure 6C-H). These data demonstrate that altered neuronal complexity could be a feature of chronic migraine, and that restoration of these changes may be a marker of effective migraine treatment.

**Figure 6.**
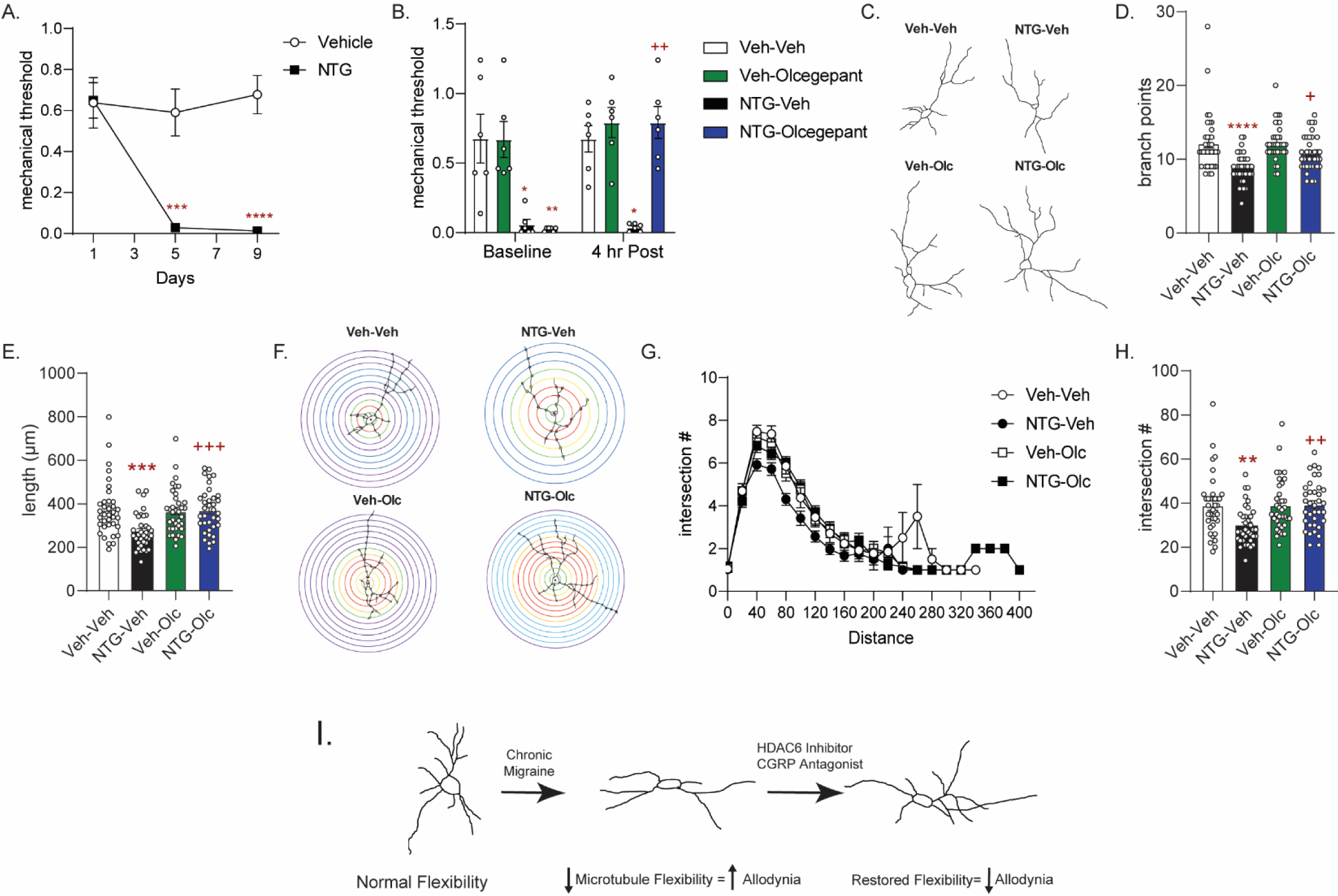
Treatment with the CGRP receptor antagonist, olcegepant, blocks NTG-induced chronic allodynia and reverses blunted cytoarchitecture. (A) Periorbital mechanical thresholds were accessed prior to Vehicle/NTG administration on days 1, 5 and 9. NTG produced cephalic allodynia; p<0.001 effect of drug, time, and interaction, two-way RM ANOVA and Holm-Sidak post hoc analysis. ***p<0.001, ****p<0.0001, relative to vehicle on same day 1 n=12/group. (B) Mice were treated with olcegepant (0.1 mg/ml, IP) or vehicle (0.9% NaCl) and tested 4 hours later. Olcegepant significantly reversed chronic cephalic allodynia. p <0.05 effect of treatment, drug, time, and interaction. Three-way ANOVA and Holm-Sidak post hoc analysis. *p<0.05, **p<0.01 relative to Vehicle-Vehicle, ++p<0.01 relative to NTG-Vehicle at the same time point n=6/group. (C) Representative neuron tracing of mice that were chronically treated with Vehicle or NTG (10 mg/kg, IP) every other day for 9 days and on day 10 were treated with olcegepant or vehicle and sacrificed 4 hours later for Golgi staining. (D) The number of branch points/neuron were significantly decreased following NTG treatment (NTG-Vehicle); while NTG-olcegepant treated mice showed increased branching compared to the NTG-Vehicle treatment alone. p<0.05 effect of chronic treatment, drug treatment, and interaction, two-way ANOVA and Holm-Sidak post hoc analysis. ****p<0.0001 NTG-Veh compared to Veh-Veh, +p<0.05 NTG-olcegepant compared to NTG-Veh. (E) NTG also decreased total length/neuron; while NTG-olcegepant treated mice showed restored length compared to the NTG-Vehicle treatment alone p<0.01 effect of NTG treatment, drug treatment, and interaction, two-way ANOVA and Holm-Sidak post hoc analysis. ***p<0.001 NTG-Veh compared to Veh-Veh, +++p<0.001 NTG-olcegepant compared to NTG-Veh. (F) Representative Sholl analysis image of neuronal complexity in the four groups. (G) Sholl analysis broken up by 20 voxel distances reveal differences between NTG-Veh treatment relative to the other groups. (H) NTG-Veh results in significantly fewer total interactions relative to the Veh-Veh treatment and NTG-olcegepant mice had significantly more total interactions compared to NTG-Veh p<0.05 effect chronic treatment, drug treatment, and interaction, two-way ANOVA and Holm-Sidak post hoc analysis. **p<0.01 NTG-Veh compared to Veh-Veh; ++p<0.01 NTG-Veh compared to NTG-olcegepant. For all analysis n=6 mice/group, 6 neurons per mouse. (I) Schematic summary of findings. Endogenously there is a balance of acetylated and deacetylated α-tubulin which regulates optimal neuronal complexity. In the case of chronic migraine, there is a disbalance resulting in decreased neuronal complexity. HDAC6 or CGRP receptor inhibition restores tubulin dynamics and neuronal complexity and correspondingly decreases chronic migraine-associated symptoms.

## Discussion

Our results indicate that in models of chronic migraine there is a dysregulation of cellular plasticity resulting in decreased neuronal complexity. We found that following the establishment of chronic cephalic allodynia in the NTG model of migraine-associated pain, there was a decrease in the number of branch points, combined neurite length, and interactions of neurons within the TNC, PAG, and somatosensory cortex. With this newly discovered phenomenon we sought to mitigate this decrease through inhibition of HDAC6 which we found to restore neuronal complexity and correspondingly, inhibit allodynia. We found that the cytoarchitectural changes were not just NTG induced but were also prominent following CSD. Reduction in neuronal complexity was also observed in this model of migraine aura, and again HDAC6 inhibition restored neuronal plasticity and decreased the number of CSD events. The latter effect is a hallmark of migraine preventive drugs. Furthermore, we found that a migraine specific treatment, CGRP receptor inhibition, also restored cytoarchitectural changes. Together our results demonstrate a novel mechanism of chronic migraine and reveal HDAC6 as a novel therapeutic target for this disorder (Figure 6I).

We used the NTG model in this study as it is a well-validated model of migraine [33]. NTG is a known human migraine trigger and has been used as a human experimental model of migraine [34]. Similar to humans, NTG produces a delayed allodynia in mice [35], as well as photophobia and altered meningeal blood flow [36, 37]. Chronic intermittent administration of NTG is used to model chronic migraine [32, 38-41]. Although, compared to humans, much higher doses of NTG are required in rodents, the allodynia induced in mice is inhibited by migraine-specific medications, such as sumatriptan [35, 38, 42] and CGRP targeting drugs [32], as well as the migraine preventives propranolol and topiramate [43, 44]. Further, mice with human migraine gene mutations are more sensitive to NTG [45]. Systemic administration of NTG also causes cellular activation throughout nociceptive pathways including in the TNC and brainstem [44, 46-48]. Correspondingly, we also observed changes in neuronal complexity in the TNC, as well as in the PAG and somatosensory cortex, regions heavily involved in pain processing. Alterations in these regions could contribute to allodynia or interictal sensitivity observed in chronic migraine patients. We did not observe any alterations in neuronal complexity in the nucleus accumbens which is commonly associated with processing of reward and motivation. RNA-Seq experiments from our lab have also showed that the nucleus accumbens shows very different responses to chronic NTG relative to parts of the trigeminovascluar system [49]. Importantly, we observed no change in neuronal cytoarchitecture in the lumbar spinal cord, a region involved in peripheral but not cephalic pain processing. These findings suggest that neuronal cytoarchitectural changes may be a common maladaptive mechanism driving migraine chronicity.

While these results are the first of their kind to demonstrate cytoarchitectural changes in models of chronic migraine, alterations in neuronal plasticity have been described previously in models of neuropathic pain. A mouse model of chronic constriction injury of the sciatic nerve reduced neurite length in GABA neurons within lamina II of the spinal cord [50]. Another group observed that following spared nerve injury, there were decreases in the number of branches and neurite length of hippocampal neurons but increases in spinal dorsal horn neurons [51]. Together these studies, along with those reported here, suggest adaptations of the central nervous system to chronic pain culminate in alteration of the neuronal cytoarchitecture.

We also observed decreased neuronal complexity in CSD, a migraine model mechanistically distinct from NTG. CSD is thought to underlie migraine aura and reflects changes in cortical excitability associated with the migraine brain state [52, 53]. Previous studies also support the idea of cytoarchitectural alterations accompanying spreading depression/depolarization events, and have mainly focused on dendritic morphology. Neuronal swelling [54] and dendritic beading [55] have been observed following spreading depression events. CSD also resulted in alterations in dendritic structure [54, 56] and volumetric changes [54]. Further, Steffensen et al. also showed decreased microtubule presence in dendrites following spreading depression in hippocampal slices, again implying alterations in cytoarchictural dynamics [55]. Microtubules have been shown to disassemble in response to increased intracellular calcium [57]; and the increased calcium influx following spreading depolarization [58] may facilitate this breakdown. Tubulin acetylation is associated with increased flexibility and stability of microtubules [11]. One way in which HDAC6 inhibitors could attenuate CSD is through increased tubulin acetylation, thus counteracting cytoarchitectural changes produced by CSD events. Furthermore, multiple reports indicate that CSD can activate the trigeminovascular complex, and evoke cephalic allodynia in rodents [59-64]. We also observed decreased neurite branching in the TNC following CSD, further linking CSD to head pain processing. Combined, these data suggest that CSD has an impact on neuronal morphology that contributes to migraine pathophysiology.

Proper acetylation of microtubules is necessary for a variety of cellular functions including appropriate neurite branching [12], cell response to injury [12], mitochondrial movement [13, 14], anchoring of kinesin for microtubule mediated transport [65] and regulation of synaptic G protein signaling [17, 66, 67]. Knockout of the α-tubulin acetylating enzyme, α-TAT1, in sensory neurons results in profound deficits in touch [68]. Furthermore, Charcot-Marie-Tooth (CMT) disease is a hereditary axonopathy that affects peripheral nerves resulting in damage to both sensory and motor function. Mouse models of CMT reveal deficits in mitochondrial transport in the dorsal root ganglia (DRG) due to reduced acetylation of α-tubulin, while HDAC6 inhibition ameliorates CMT-associated symptoms [69, 70]. HDAC6 inhibitors were also found to be effective in models of chemotherapy induced neuropathic pain [15, 71]. In mice, HDAC6 inhibitors effectively reduced chemotherapy-induced allodynia following treatment with vincristine [15] or cisplatin [71]. In addition, both groups found that chemotherapy blunted mitochondrial transport in sensory neurons, an effect that was restored by HDAC6 inhibition. A recent study also found that HDAC6 inhibition was effective in models of inflammatory and neuropathic pain [72]. Together with our study, these data further demonstrate the importance of HDAC6 in regulating acetylated α-tubulin and how tubulin acetylation state can regulate sensory dysfunction; thus supporting the potential of HDAC6 inhibitors for the treatment for chronic pain conditions.

We observed that chronic migraine resulted in decreased neuronal complexity and therefore focused on the role of HDAC6 in tubulin acetylation and cytoarchitectural dynamics [10, 12]. However, HDAC6 also regulates Hsp90 and cortactin [7]. HDAC6 deacetylates Hsp90, which plays an important role in glucocorticoid receptor maturation and adaptation to stress [73]. A previous study using a model of social defeat stress showed that HDAC6 knockout or inhibition decreased Hsp90-glucocorticoid receptor interaction and subsequent glucocorticoid signaling, thus encouraging resilience [27]. In line with these findings, HDAC6 inhibitors also show antidepressant-like effects [16, 25], and membrane-associated acetylated tubulin is decreased in humans with depression [67]. Changes in neuronal complexity may serve as a common mechanism underlying the chronicity of psychiatric and neurological diseases. Further, HDAC6 also directly deacetylates cortactin, a protein that regulates actin dependent cell motility [74]. Future studies will focus on identifying the precise mechanism by which HDAC6 impacts neuronal complexity in the chronic migraine state.

We investigated whether a current migraine treatment strategy, CGRP receptor inhibition, could ameliorate the cytoarchitectural changes induced by chronic migraine-associated pain. We found a good correlation between the anti-allodynic effects of olcegepant and its ability to restore neuronal complexity in the chronic NTG model. In contrast to ACY-738, olcegepant had no effect on vehicle treated mice and only recovered but did not increase neuronal branching, length, or intersections in the NTG treated group. These results, along with the finding that NTG did not alter complexity in the lumbar spinal cord or nucleus accumbens, help to confirm that the changes in neuronal cytoarchitecture following NTG are associated with migraine mechanisms. In addition, this study suggests a possible mechanism in which recovered neuronal complexity is a marker of effective migraine medication. Our findings also open up research on the signaling mechanisms that link migraine relief with tubulin dynamics. One possibility is that decreased neuronal cytoarchitecture is driven through a CGRP related mechanism and direct antagonism of the CGRP receptor reverses these alterations. NTG treatment was previously found to upregulate CGRP in the blood, TNC, and dura mater demonstrating a direct link between NTG and CGRP [75-77]. Future studies will explore the relationship between CGRP and other migraine therapies with neuronal cytoarchitecture.

Our results reveal a novel cytoarchitectural mechanism underlying chronic migraine, and imply that this disorder may result from attenuation of neurite outgrowth and branching. This blunted complexity would consequently decrease neurnoal plasiticity and the ability of the cell to respond to its environment. We propose that this decreased flexibility could also encourage the strengthening of maladaptive synapses and prevent the formation of potentially beneficial synapses. Human imagining studies reveal decreased cortical thickness [78], and grey matter reductions in the insula, anterior cingulate cortex, and amygdala of migraine patients [79]. Interestingly, a significant correlation was observed between gray matter reduction in anterior cingulate cortex and frequency of migraine attacks [79]. These structual changes could reflect decreased neuronal complextity in combination with other factors.

We propose that strategies targeted towards pathways regulating neuronal cytoarchicture may be an effective approach for the treatment of chronic migraine. Although HDAC6 inhibition increased neuronal complexity in vehicle or sham animals, this did not affect normal pain processing or cortical excitability, respectively. Additionally, constitutive knockout of HDAC6 produces viable offspring with few phenotypic changes other than resilience to stress [80]. Together these results suggest that HDAC6 inhbitors may restore cellular adaptations induced by chronic disease states but may not otherwise affect healthy physiological function. Chronic migraine may result from attenuation of neurite outgrowth and branching and compounds that reverse this adaptation, such as HDAC6 inhibitors, may contribute to the migraine therapeutic armamentarium.

## Materials and Methods

### Animals

Experiments were performed on adult male and female C57BL6/J mice (Jackson Laboratories, Bar Harbor, ME. USA) weighing 20-30g. Mice were group housed in a 12h-12h light-dark cycle, where the lights were turned on at 07:00 and turned off at 19:00. Food and water were available ad libitum. All experiments were conducted in a blinded fashion by 1-3 experimenters. Weight was recorded on each test day for all experiments. All experimental procedures were approved by the University of Illinois at Chicago Office of Animal Care and Institutional Biosafety Committee, in accordance with Association for Assessment and Accreditation of Laboratory Animal Care International (AAALAC) guidelines and the Animal Care Policies of the University of Illinois at Chicago. All results are reported according to Animal Research: reporting of In Vivo Experiments (ARRIVE) guidelines. No adverse effects were observed during these studies, and all animals were included in statistical analysis.

### Sensory Sensitivity testing

Different groups of animals were used for each experiment. Mice were counter-balanced into groups following the first basal test for mechanical thresholds. Mice were tested in a behavior room, separate from the vivarium, with low light (∼35-50 lux) and low-noise conditions, between 09:00 and 16:00. Mice were habituated to the testing racks for 2 days before the initial test day, and on each subsequent test days were habituated for 20 min before the first test measurement. For cephalic measures mice were tested in 4 oz paper cups. The periorbital region caudal to the eyes and near the midline was tested. For experiments testing peripheral mechanical responses, the intraplantar region of the hindpaw was assessed. Testing of mechanical thresholds to punctate mechanical stimuli was tested using the up-and-down method.

The selected region of interest was stimulated using a series of manual von Frey hair filaments (bending force ranging from 0.008 g to 2 g). A response of the head was defined as shaking, repeated pawing, or cowering away from the filament. In the hind paw a response was lifting of the paw, shaking, or licking the paw after stimulation. The first filament used was 0.4 g. If there was no response a heavier filament (up) was used, and if there was a response a lighter filament (down) was tested. The up-down pattern persisted for 4 filaments after the first response.

### Nitroglycerin model of chronic migraine

Nitroglycerin (NTG) was purchased at a concentration of 5 mg/ml, in 30% alcohol, 30% propylene glycol and water (American Reagent, NY, USA). NTG was diluted on each test day in 0.9% saline to a concentration of 1 mg/ml for a dose of 10 mg/kg. Mice were administered NTG or vehicle every other day for 9 days. Animals used in cephalic experiments were tested on days 1, 5, and 9. On test days a basal threshold was measured then animals were treated with either NTG or vehicle and then put back in the testing racks and subsequently tested 2 h later for the post-treatment effect.

### Cortical Spreading Depression Model

The procedure for the cortical spreading depression (CSD) model is based on work previously published by Ayata[81] that is commonly used to screen potential migraine preventatives and further used in our own work Pradhan and colleagues[42]. Mice were grouped into sham and CSD groups and then further subdivide into ACY-738 (50 mg/kg, IP) or vehicle (i.e., Sham-ACY, Sham-Veh, CSD-ACY, CSD-Veh). To make the thinned skull cortical window, mice were anesthetized with isoflurane (induction 3-4%; maintenance 0.75 to 1.25%; in 67% N_2_ / 33% O_2_) and placed in a stereotaxic frame on a homoeothermic heating pad. Core temperature (37.0±0.5°C), non-peripheral oxygen saturation (∼ 99%), heart rate, and respiratory rate (80–120 bpm) were continuously monitored (PhysioSuite; Kent Scientific Instruments, Torrington, CT, USA). Mice were frequently tested for tail and hind paw reactivity to ensure that the anesthesia plane was maintained.

To verify CSD events, optical intrinsic imaging (OIS) and electrophysiological recordings were performed as previously described[42]. Briefly, following anesthesia, the skin from the skull was detached and a rectangular region of ∼2.5 x 3.3 mm^2^ (∼0.5 mm from sagittal, and ∼1.4 from coronal and lambdoid sutures) of the right parietal bone was thinned to transparency with a dental drill (Fine Science Tools, Inc., Foster City, CA, USA). Mineral oil application improved transparency of cortical surface parenchyma and vasculature for video recording. A green LED (530 nm) illuminated the skull throughout the experiment (1-UP; LED Supply, Randolph, VT, USA). Cortical surface reflectance detected by OIS was collected with a lens (HR Plan Apo 0.5×WD 136) through a 515LP emission filter on a Nikon SMZ 1500 stereomicroscope (Nikon Instruments, Melville, NY, USA). Images were acquired at 1–5Hz using a high-sensitivity USB monochrome CCD (CCE-B013-U; Mightex, Pleasanton, CA, USA) with 4.65-micron square pixels and 1392×1040 pixel resolution.

Lateral to the thinned window two burr holes were drilled around the midpoint of the rectangle. These burr holes were deeper than the previously drilled skull region such that the dura was exposed but not broken. To record local field potentials (LFPs) an electrode (in a pulled glass pipette filled with saline) was inserted into one burr hole and attached to an amplifier. A separate ground wire, placed underneath the skin caudal to the skull, grounded this set up and LFPs were recorded for an hour to ensure a stable baseline and recovery from any surgically induced CSDs. After an hour of stabilization, a second pulled glass pipette was filled with 1 M KCl and placed into the more rostral burr hole, avoiding contact with the brain or the surrounding skull. An initial flow of KCl was pushed to begin and then an even flow was held so that there was a constant small pool of KCl that filled the burr hole. Excess liquid was removed with tissue paper applied next to the burr hole. Regardless of grouping the CSD recording continued for 3600s after the initial drip of KCl. Mice were euthanized by anesthetic overdose followed by decapitation.

### Golgi Staining

Golgi staining was performed according to the FD Rapid Golgi Stain kit (FD Neurotechnologies). For NTG or Veh treated mice, they underwent the chronic NTG model and on day 10, 4 h after ACY-738 treatment or vehicle, mice were anesthetized with isoflurane and then euthanized. After euthanizing the brains were removed rapidly. The tissue was then rinsed briefly in double distilled water. Tissue was then placed in the impregnation solution that was an equal amount of solutions A and B that was prepared at least 24 h in advance. After the first 24 h the brain was placed in new impregnation solution and then stored for 1 week in the dark. The brains were then transferred to solution C, which was also replaced after the first 24 h. After replacing solution C the brains were stored at room temperature for 72 h more. Following solution C, brains were flash frozen in 2-methyl butane and cryostat cut at −20°C into 100 μm slices. The slices were mounted onto gelatin coated slides and secured by a drop of solution C placed onto each slice. These slides were then left to dry naturally in the dark.

### Neurite Tracing

After processing images were taken at 20x magnification and a Z-stack was created based on different levels of focal plane. After the Z-stack was created the FIJI program Simple Neurite Tracer was used to trace the processes of the neuron.

Furthermore, after tracing the neurons were analyzed using Simple Neurite Tracer [82] software to assess the number of branch points from each neuron, overall length of the neuron, and Sholl Analysis. Sholl Analysis was performed by placing a center ROI point at the center of the soma and producing consecutive circles every 20 pixels for the entire body of the neuron. Intersections were counted based on the number of times a neurite crossed each of these consecutive circles. These data were compiled per neuron and then brought into one Masterfile.

### Neuron Selection

Throughout tracing all tracers were blinded to which group the images belonged to. For all brain regions analyzed, six to eight relatively isolated neurons were randomly chosen per mouse. The selected neurons were fully impregnated with Golgi stain and relatively complete. An atlas was used along with clear anatomical markers to ensure the neurons were being taken from their described region of interest. Neurons characterized for the trigeminal nucleus caudalis region were taken only from the outer lamina of caudal sections. Neurons analyzed for somatosensory cortex were all taken from layer IV of the primary somatosensory barrel cortex. To ensure a homogenous cell population only pyramidal cells were selected. The most complex neurons were chosen for analysis in all regions. Previously, it was shown that dendritic complexity was directly correlated to soma size. To ensure that the NTG group where not just smaller in size we directly compared soma diameter of neurons in the NTG and Veh group. There was no significant difference in soma size between these two groups (Veh 9.258 ± .2515 and NTG 9.192 ± .2782, student run t-test p= 0.8608). Two individuals traced all cells. Interrater reliability was determined by having each tracer trace 5 neurons in their entirety. Pearson product correlations were accessed in three measures; number of branches, total dendritic length, and total intersection number through Sholl analysis and found to be 0.91, 0.94, and 0.95 respectively. All tracings of neurons were re-examined by the primary tracer (Z.B.) to assure quality control.

### Drug Injections

All injections were administered at 10 ml/kg volume, intraperitoneally (IP), unless otherwise indicated. ACY-738 was dissolved in a 5% DMSO saline solution, which was used as the vehicle control. RN-73 was dissolved in 10% DMSO, 10% Tween-80, and 80% saline and was injected 1 mg/kg or 10 mg/kg, this mixture was also used as the vehicle control group. ASV-85 was dissolved in 15% DMSO, 15% Tween-80, and then 70% saline, this mixture was also used as the vehicle control group. ASV-85 was injected at 1 mg/kg dose. TSA was dissolved in 20% DMSO solution in 80% 0.01M PBS and injected at a dose of 2 mg/kg, this same solution was used for the vehicle. Olcegepant was dissolved in saline solution and was injected at a 0.1 mg/kg dose. For the CSD experiments ACY-738 was injected 3 hours before starting the surgery so that it would reach its peak efficiency of 4 h by the time the CSD event started.

### Quantitative RT-PCR

Total RNA was isolated from flash frozen brain punches using the RNeasy Plus Mini kit from Quiagen. RNA samples were reverse transcribed to single-stranded cDNA. cDNA transcription was used following the protocol from Superscript III (Life Technologies) and the TaqMan Gene Expression Assay system (Applied Biosystems). Glyceraldehyde-3-phosphate dehydrogenase (GAPDH, Hs02758991_g1) was used as a housekeeping gene. The threshold cycle (CT) of each target product was determined and CT values between HDAC6 transcripts and housekeeping genes were calculated (ΔCT). The fold change (2^- ΔΔCT^) for each was calculated relative to the median ΔCT from the saline control animals.

### Immunohistochemistry

Mice were anesthetized with Somnasol (100 μl/mouse; 390 mg/mL pentobarbital sodium; Henry Schein) and perfused intracardially with 15 ml of ice-cold phosphate-buffered saline (0.1 M PBS, pH 7.2) and subsequently 50 mL of ice-cold 4% paraformaldehyde (PFA) in 0.1M PBS (pH 7.4). Whole brain and trigeminal ganglia (TG) were harvested and overnight left to post-fix in 4% PFA/0.1M PBS at 4°C. Brain and TG were then cryoprotected in 30% sucrose in 0.1M PBS until they sunk. Brains were then flash frozen using 2-methyl butane over dry ice. Coronal sections of the trigeminal nucleus caudalis (TNC) and the somatosensory cortex were sliced at 20 μM and TG at 16 μM. All slices were immediately mounted onto slides after slicing in the cryostat. Slides were washed with PBST. Then a blocking solution with 5% normal donkey serum with PBST for 1 h at room temperature. Slides were then incubated overnight at RT with the primary rabbit anti-HDAC6 antibody (1:500, courtesy of Tso-Pang Yao at Duke University) diluted in 1% NDSDT. Slides were subsequently washed with 1% NDST and then the secondary antibody was added for 2 hours at room temperature (donkey anti rabbit IgG, 1:2000). Slides were washed with 0.1 M phosphate buffer, and cover slipped with Mowiol-DAPI mounting medium. Images were taken by in a blinded manner using the EVOS FL Auto Cell Imaging system, using a 40 x objective.

### Western Blots

Samples were taken from chronically treated NTG or Vehicle mice, which received an injection of ACY-738 or Vehicle on day 10. Samples were collected 4h post-ACY/VEH. Protein concentrations were assessed using a Nanodrop 2000c spectrophotometer and equal quantities were loaded onto each Stain-Free acrylamide gel for SDS-PAGE (Bio-Rad, Hercules, CA, USA). The gels were subsequently transferred to Nitrocellulose membranes (Bio-Rad, Hercules, CA USA) for western blotting. The membranes were blocked with 5% non-fat dry milk diluted in TBS-T (10 mM Tris-HCl, 159 mM NaCl, and 0.1% Tween 20, pH 7.4) for 1 h. Following the blocking step, membranes were washed with Tris-buffered saline/Tween 20 and then incubated with an anti-acetyl-α-tubulin antibody (Lysine-40) (Sigma Clone 6-11B1), α-tubulin (Sigma), overnight at 4 °C. Membranes were washed with TBS-T and incubated with a secondary antibody [HRP-linked anti-mouse antibody IgG F(ab′)2 or HRP-linked anti-rabbit antibody IgG F(ab′)2] (Jackson ImmunoResearch, West Grove, PA, USA, catalog #115-036-072 for mouse, and catalog #111-036-047 for rabbit, RRID) for 1 h at room temperature, washed, and developed using ECL Luminata Forte chemiluminescent reagent (Millipore, Billerica, MA, USA). Blots were imaged using a Chemidoc computerized densitometer (Bio-Rad, Hercules, CA, USA) and quantified by ImageLab 3.0 software (Bio-Rad, Hercules, CA, USA). In all experiments, the original gels were visualized using BioRad stainfree technology to verify protein loading.

### Statistical Analysis

Sample size was calculated by power analysis and previous experience. Since we investigated changes at the cellular level, an individual neuron represented a single sample [27, 83]. Data analysis was performed using GraphPad Prism version 8.00 (GraphPad, San Diego, CA). The level of significance (α) for all tests was set to 0.05. Post hoc analysis was conducted using Holm-Sidak post hoc test to correct for multiple comparisons. Post hoc analysis was only performed when F values achieved p < 0.05. All values in the text are reported as mean ± SEM.

### Data availability

All data are available upon reasonable request.

## Acknowledgments

This work was funded by NIH grants: NS109862 (AAP), DA040688 (AAP), AT009169 (MMR), VA grant VA BX00149 (MMR), an Amgen Competitive Migraine Grant (AAP), UICentre for Drug Discovery (PP, AAP). MMR is a VA Research Career Scientist. ZB is a member of the UIC Graduate Program in Neuroscience. We thank Tso-Pang Yao from Duke University for the HDAC6-selective antibody.

## Author Contributions

ZB, HS, PP, SMB, MMR, AAP conceived and planned experiments, and edited the manuscript. ZB, HS, ID, KS, AT, WW, ZS, PS, CC performed experiments. VP, BK, PP synthesized compounds. ZB, HS, ID, PP, MMR, AAP analyzed data. ZB, MMR, AAP wrote the manuscript.

## Supplementary Materials

**Supplementary Figure 1.**
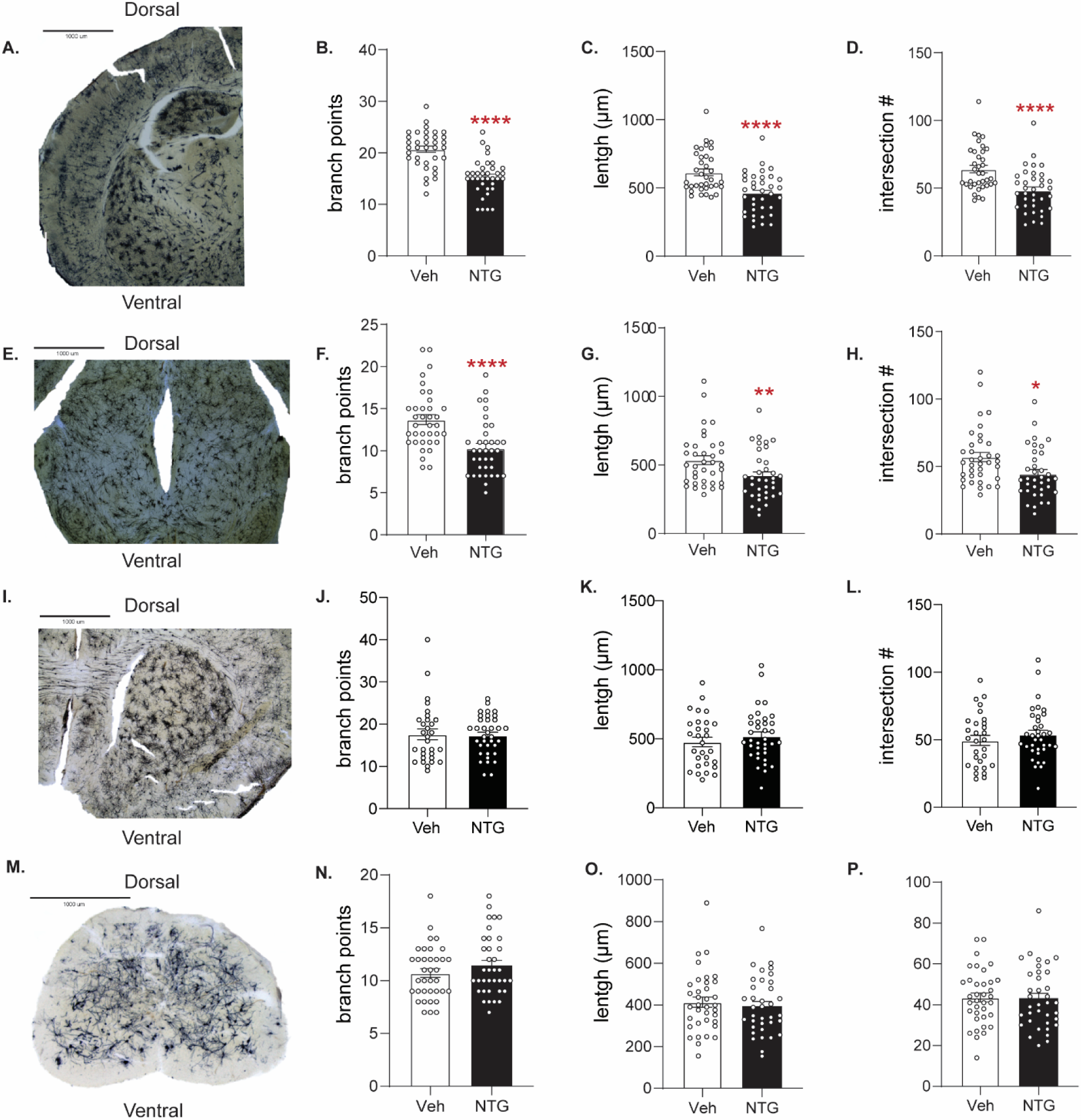
Chronic NTG treatment causes cytoarchitectural changes in the Somatosensory Cortex (SCx) and Periaqueductal Grey (PAG) but not the Nucleus Accumbens (Nac) or Lumbar Spinal Cord (LSC). Mice were treated chronic intermittently with either NTG or Vehicle for 9 days and on day 10 tissue was collected for Golgi staining. (A,E,I,M) Representative image showing Golgi staining in the SCx, PAG, Nac, and LSC. (B,F,J,N) Neurons were analyzed for number of branch points, (C,G,K,O) combined neuronal length and (D,H,L,P) total number of intersections using Sholl analysis. In the SCx (A-D) and PAG (E-H). chronic NTG resulted in significantly fewer branches, neurite length, and interactions compared to vehicle treated controls; unpaired t-test. *p<0.05, ** p<0.01, ****p<.0001. In the Nac (I-L) and LSC (M-P), chronic NTG did not produce significant changes in the number of branch points, neurite length, or interactions compared to vehicle controls. n=6 mice/group, 6 neurons/mouse.

**Supplementary Figure 2.**
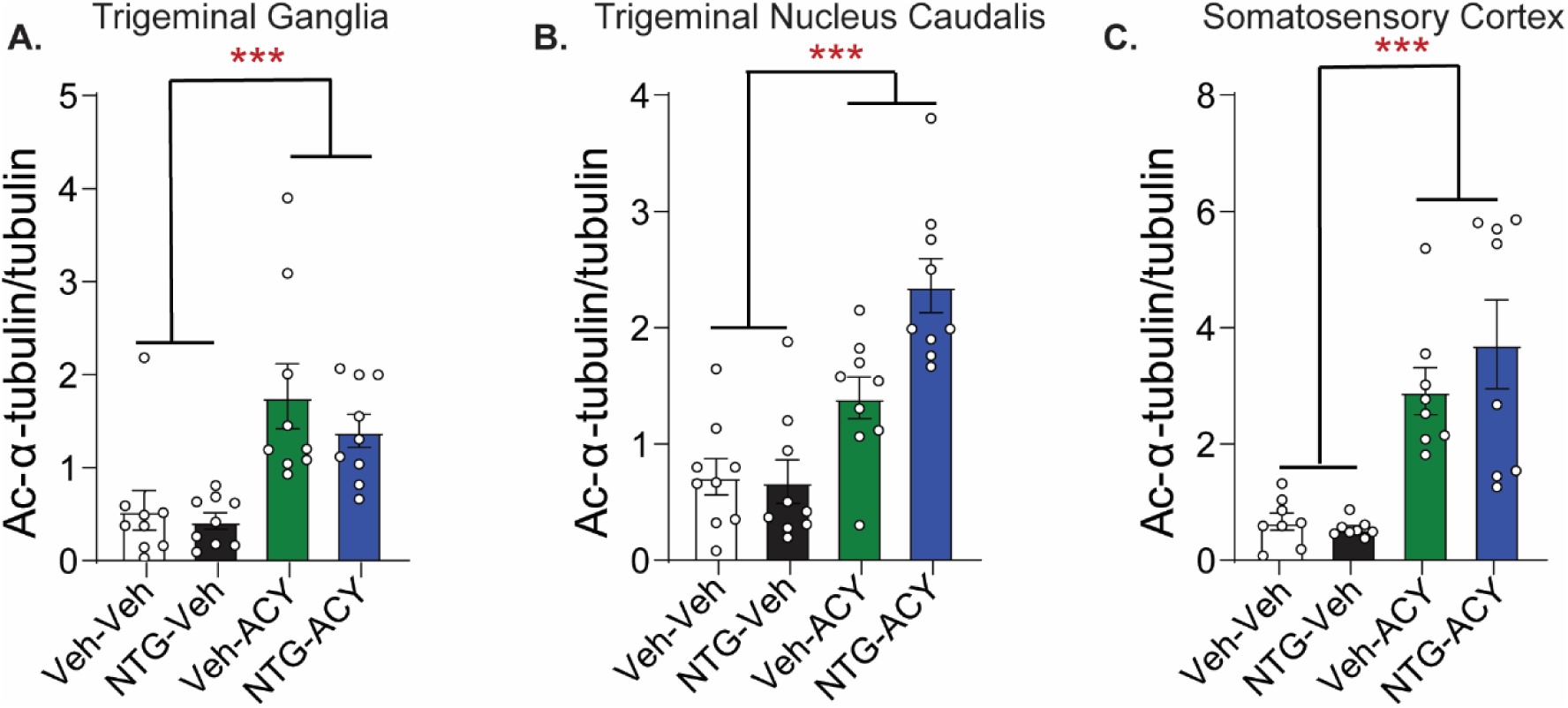
ACY-738 increased levels of acetylated α-tubulin. Mice were chronically treated with NTG or vehicle for 9 days. On day 10 mice were injected with ACY-738 (50 mg/kg IP) or vehicle (5% DMSO, 0.9% NaCl, IP) and tissue was collected 4h later. (A) Trigeminal ganglia (B) somatosensory cortex, and (C) trigeminal nucleus caudalis of mice treated with ACY-738 showed a significant increase in the ratio of acetylated α-tubulin/total α-tubulin compared to the vehicle treated mice ***p<.0001 n=8-9/group effect drug treatment two-way ANOVA.

**Supplementary Figure 3.**
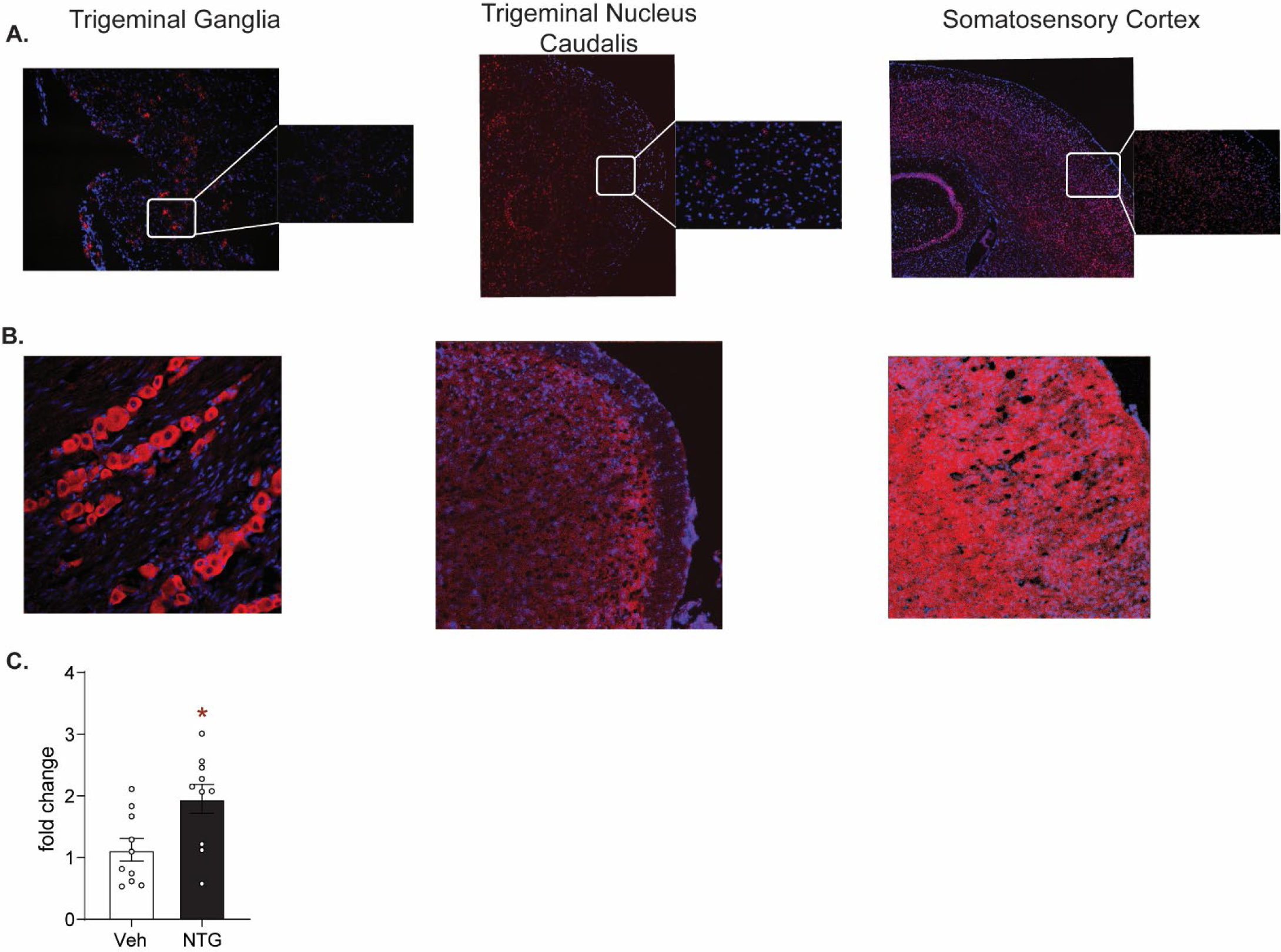
HDAC6 is expressed in migraine-processing regions and is dynamically regulated. (A.) In situ hybridization by RNAScope reveals abundant HDAC6 transcripts (red) in trigeminal ganglia, trigeminal nucleus caudalis, and somatosensory cortex (B.) Immunohistochemical staining also reveals HDAC6 expression in these regions. (C.) Mice were treated chronically with vehicle or NTG for 9 days and tissue was analyzed for HDAC6 gene expression on day 10. HDAC6 transcript levels were significantly increased following NTG treatment in the TG, unpaired t-test, *p<0.05, n=10 mice/group.

**Supplementary Figure 4.**
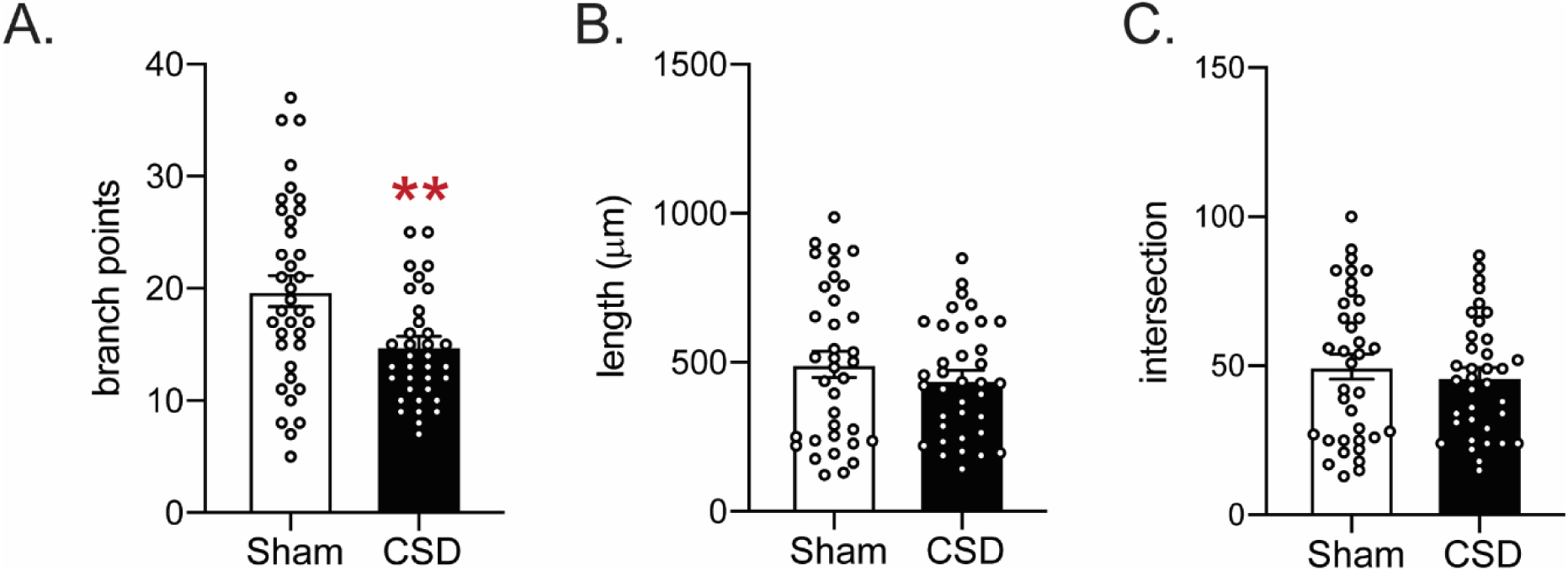
Cortical spreading depression (CSD) results in blunted neuronal complexity in the trigeminal nucleus caudalis (TNC). Mice underwent the previously described CSD procedure or a sham surgery in which no KCl was dripped onto the dura. Mice were sacrificed immediately after an hour of recording. (A) Neurons were analyzed for number of branch points, and CSD resulted in significantly fewer branch points relative to sham controls. (B) Neurons were further analyzed for combined neuronal length and no significant difference was observed. (C) Total number of intersections were analyzed using Sholl analysis and there was no significant difference. Unpaired t-test ** p<0.01 n=6/group n=6 mice/group

